# Genetic Regulation of Phenotypic Plasticity and Canalisation in Yeast Growth

**DOI:** 10.1101/066175

**Authors:** Anupama Yadav, Kaustubh Dhole, Himanshu Sinha

## Abstract

The ability of a genotype to show diverse phenotypes in different environments is called phenotypic plasticity. Phenotypic plasticity helps populations to evade extinctions in novel environments, facilitates adaptation and fuels evolution. However, most studies focus on understanding the genetic basis of phenotypic regulation in specific environments. As a result, while it’s evolutionary relevance is well established, genetic mechanisms regulating phenotypic plasticity and their overlap with the environment specific regulators is not well understood. *Saccharomyces cerevisiae* is highly sensitive to the environment, which acts as not just external stimulus but also as signalling cue for this unicellular, sessile organism. We used a previously published dataset of a biparental yeast population grown in 34 diverse environments and mapped genetic loci regulating variation in phenotypic plasticity, plasticity QTL, and compared them with environment-specific QTL. Plasticity QTL is one whose one allele exhibits high plasticity whereas the other shows a relatively canalised behaviour. We mapped phenotypic plasticity using two parameters – environmental variance, an environmental order-independent parameter and reaction norm (slope), an environmental order-dependent parameter. Our results show a partial overlap between pleiotropic QTL and plasticity QTL such that while some plasticity QTL are also pleiotropic, others have a significant effect on phenotypic plasticity without being significant in any environment independently. Furthermore, while some plasticity QTL are revealed only in specific environmental orders, we identify large effect plasticity QTL, which are order-independent such that whatever the order of the environments, one allele is always plastic and the other is canalised. Finally, we show that the environments can be divided into two categories based on the phenotypic diversity of the population within them and the two categories have differential regulators of phenotypic plasticity. Our results highlight the importance of identifying genetic regulators of phenotypic plasticity to comprehensively understand the genotype-phenotype map.

## INTRODUCTION

A single genotype cannot have high fitness in all conditions. Instead different genotypes show varying degrees of fitness in different environments, and therefore phenotype of a genotype is dependent on the environment. The ability of a single genotype to show different phenotypes in different environments is called phenotypic plasticity [1]. On the other hand, ability of a genotype to show the same phenotype independent of the environment is termed as canalisation [2]. Phenotypic plasticity facilitates adaptation to novel environments by allowing the population to exhibit a diverse range of phenotypes [3]. It is ubiquitous in nature and shown to be a major force in adaptation, be it adaptation to climate change, altitude, nutrition, multi-cellularity, etc. [4,5]. Consequently, phenotypic plasticity is one of the major drivers of evolution [2,6,7].

During adaptation, stabilising selection acts on the population such that the phenotype gets stabilised or canalised within an environment and across multiple environments [8]. One of the ways, this canalisation is proposed to get perturbed is when this adapted population encounters a novel or rare environment. This perturbation of canalisation allows the population to exhibit a range of phenotypes thus facilitating adaptation. Canalisation and plasticity are dynamic, mutually dependent processes and a population switches between these two states depending on the environments encountered [9,10]. While a canalised phenotype would be beneficial in environments to which the population has adapted to, a plastic phenotype would be advantageous in a novel or rare environment [6]. Hence the same genotype is capable of showing a canalised or plastic behaviour depending on the environments considered and different genetic regulators may regulate phenotypic plasticity in varying environments.

While the importance of plasticity in adaptation and evolution has been established by multiple studies [11], these studies are mostly conducted in naturally occurring populations. Therefore, while evidence for phenotypic plasticity has been documented in multiple organisms across diverse phenotypes, its genetic regulation is not clearly understood. Additionally, most studies that attempt to understand the genetic regulation of a phenotype focus on either a single environment or multiple environments independently [12-14]. As a result, while our knowledge about genetic regulation of a phenotype in different environments is fairly comprehensive, we do not understand the genetic regulation of plasticity and canalisation across diverse environments. While phenotypic plasticity is mainly invoked to study the adaptability of natural populations, its ubiquity and role in evolution indicates that it should also be important for understanding the genetic architecture of complex traits [15].

Quantitative trait locus (QTL) mapping provides a good way to identify regulators of phenotypic plasticity. Phenotypes of most loci show environment dependence [16]. By this definition, all loci showing gene-environment interaction (GEI) exhibit phenotypic plasticity. However, a plasticity QTL is a locus whose one allele shows a canalised behaviour whereas the other allele shows phenotypic plasticity across diverse environments [17] (Fig 1A, 1B). If two genetically diverse strains have encountered and adapted to varied environments, or adapted to the same environments using different mechanisms, then crossing these strains will disrupt these mechanisms and allow identification of loci with differential plasticity in this biparental population.

**Fig 1:**
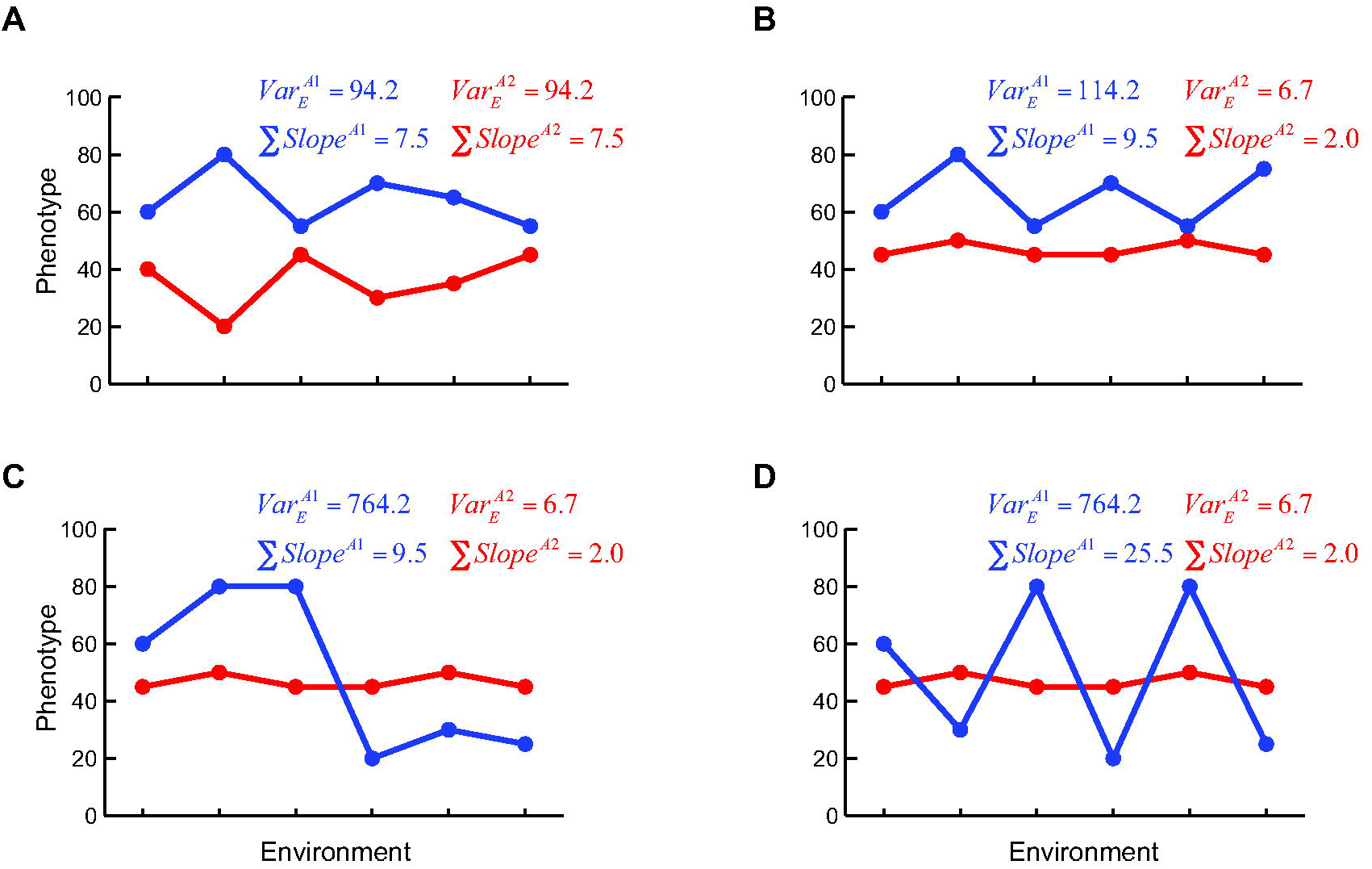
Schematic showing dependence of phenotypic plasticity parameters on the order of the environments. Genotype A1 and A2 are represented in blue and red colours respectively. *Var_E_* refers to environmental variance whereas ∑*Slope* refers to sum of slopes, as described in Methods. y-axis denotes the phenotype and x-axis denotes discrete environments arranged in different orders. (A) Genotype A1 and A2 have significant differences in multiple environments but are both equally plastic. (B) A1 is plastic and A2 is canalised. (C) and (D) shows the same environments arranged in different orders which have no effect on environmental variance but have different impact on reaction norms or sum of slopes.

While multiple studies have performed QTL mapping to identify plasticity QTL, they were either done across pairs of environments or continuums of environments [17,18]. However, in nature, populations encounter diverse environments, capable of affecting the phenotype, either simultaneously or consecutively. In-lab evolution studies have shown that the order of environments encountered during the course of evolution can dictate which alleles eventually get fixed in a population [19]. Parallel to this, it is probable that the order of encountering these environments would determine the plasticity of the genotype, which would in turn determine the selection forces that act on it (Fig 1). Different genotypes can show different ranges of phenotypic plasticity depending on the order of the environments and different parameters are required to capture the plasticity in different environmental groups (Fig 1C, 1D). Hence in order to comprehensively identify the regulators of phenotypic plasticity, both diversity of environments and their order should be considered.

In this paper, we asked the following questions: can we identify plasticity QTL across a large number of heterogeneous environments? How do these plasticity QTL respond to different types and orders of environments? Finally, what is the association between pleiotropic regulators of the phenotype and plasticity regulators? Are loci that regulate plasticity and that are pleiotropic across multiple environments same such that all pleiotropic loci contribute to plasticity, or are these loci different and hence not identified in environment-specific QTL mapping?

*S. cerevisiae* provides an ideal system to identify the genetic regulators of phenotypic plasticity, since environment serves as both external stimulus as well as signalling cue for this unicellular, sessile, organism. Yeast growth is highly responsive to environments and has been shown to be differentially regulated in different environments [16,20,21]. In this study, using growth phenotype measured in 34 diverse environments for a large yeast biparental population [13], we measured phenotypic plasticity using two statistics: an environmental order-independent statistic – *Environmental variance* (*Var_E_*), and an environmental order-dependent statistic, *Sum of slopes* (reaction norms) (∑*Slope*) (Fig 1). Fig 1 shows that both these parameters capture different aspects of phenotypic plasticity. Fig 1A shows that genotypes with difference in phenotype across diverse environments do not necessarily have differential plasticity; 1B shows two genotypes with differential plasticity; and Fig 1C and 1D show that while the environmental order has no bearing on environmental variance, the value of the reactions norms is highly sensitive to the order of the environments encountered. We use these two parameters to identify loci with differential effects on phenotypic plasticity, plasticity QTL. To the best of our knowledge, our study is the first study to identify genetic regulation of phenotypic plasticity and canalisation across such a diverse set of environments. These genetic regulators of phenotypic plasticity may play an important role in explaining missing heritability and understanding the genetic regulation of complex traits especially human disease that are influenced by multiple environmental conditions.

## METHODS

### Dataset

The raw growth data used in this study was derived from a previously published study by Bloom et al. [13], in which the experimental procedures are described in detail. The data we used was generated for 1,008 segregants derived from a cross between yeast strains BY (a laboratory strain) and RM11-1a (a wine isolate, indicated as RM). These segregants were genotyped for a total of 11,623 polymorphic markers and were grown and phenotyped for colony size in 46 different conditions. Of these 46 conditions, we selected 34 conditions based on following three criteria: (i) segregant phenotype in a particular environment should show normal distribution; (ii) environments should be closer or mimic naturally occurring environmental conditions; (iii) since different degrees of environmental stresses can invoke correlated phenotypes, biasing our analysis, only heterogeneous environments were chosen. This filtering removed environments like high temperature growth (37°C), rapamycin, pH and temperature gradients, etc.

### Single QTL Mapping

QTL mapping was carried out as described previously [21]. In brief, the R/qtl package [22,23] was used to identify QTL separately for colony size in each environment. QTL were identified using the LOD score, which is the log_10_ of the ratio of the likelihood of the experimental hypothesis to the likelihood of the null hypothesis [23]. An interval mapping method (‘scanone’ function in R/qtl) was used to compute this LOD score using the Haley-Knott regression algorithm [22].

The following formula was used to calculate the F-score, which was further used to derive the LOD score. At a particular marker, let segregant *i*’s phenotypic value be *y_ij_* where *j* can take two values (*j* = 1: BY allele and *j* = 2: RM allele).

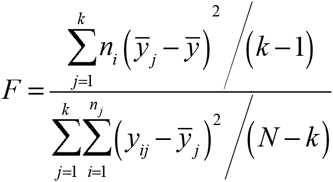

here, *N* is the total number of segregants, *n_1_* and *n_2_* are the number of segregants having the BY and RM allele respectively (*k* = 2) and *y_i_* is the genotypic mean of allele *j*.

Let *df* denote the degrees of freedom (*df* = 1 for a backcross and *df* = 2 for an intercross). The LOD score is accordingly derived as follows:

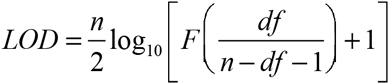

Under the null hypothesis, there is no significant difference in the means at the marker under consideration while under the alternative hypothesis, there is a presence of a QTL.

### Plasticity QTL Mapping

Plasticity QTL mapping was performed using the same methodology as described for QTL mapping, using environmental variance and sum of slopes as phenotypes, instead of colony size.

Environmental variance (*Var_E_*) was computed for each segregant separately for high (*Hv*) and low (*Lv*) variance environments:

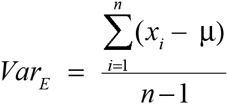

where, *x* is phenotype of a segregant in an environment, μ is the average phenotype across *n* environments. *n* = 10 for *Hv* and *n* = 24 for *Lv* environments. For mapping in sub-groups of *Hv* environments, *n* was 3 and 4, respectively.

Sum of slopes (∑*Slope*) was calculated for each segregant for each order of environments using the following formula:

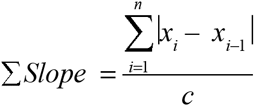

Where *n* is number of environments in a given order, *x* is the phenotype in the environment and *c* is the constant that represents difference between the two environments. Since all the environments are heterogeneous discrete environments and do not represent a continuum, the difference between them is always a constant, thus *c* was given a value of 1.

### Random orders and allele specific plasticity QTL

Environmental order for calculating the sum of slopes was determined in two different ways: random orders, where for both *Hv* and *Lv* environments independently, 10 random orders of environments were generated. For a particular order, each environment was given a single unique position, such that there were no repetitions of environments. Sum of slopes was calculated for each segregant for each order and QTL mapping was done for each order separately. Allele specific orders separately for both BY and RM alleles and for both *Hv* and *Lv* environments independently, the environments were ordered such that the mean of the segregants carrying a particular allele have the least possible sum of slopes. In other words, the mean of the population is canalised across the environmental order. Sum of slopes was calculated for this order for all segregants and QTL mapping was performed.

## RESULTS

### Environments fall into two categories based on the variance of the segregants

In the previously published dataset [13], we computed the variance of all segregants across 34 environments to identify the range of phenotypic plasticity exhibited by the individuals of the population. A higher variance would indicate high diversity of the phenotype of the segregant across the environments (high phenotypic plasticity) whereas a low variance would suggest similar phenotype across all environments (canalisation). The phenotypic variance showed a normal distribution indicating that it was a complex trait with a fraction of individuals showing highly canalised and highly plastic behaviour (Fig 2A, S1A, S1B). There was no association between the variance and average phenotype of the segregants (*R^2^* = 0.0007) indicating that segregants with both high and low average phenotype could show high variance.

**Fig 2:**
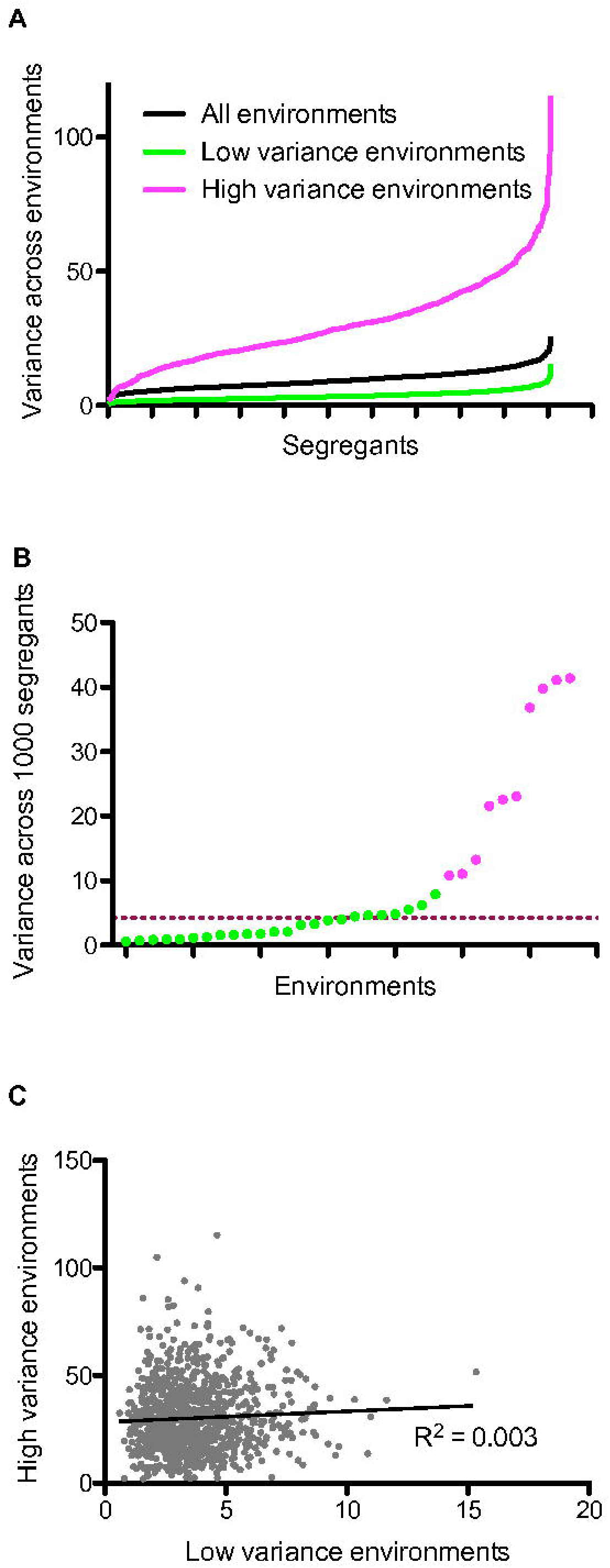
Categorisation of environments based on phenotypic variance. (A) Phenotypic variance of ~1000 segregants (x-axis) across different environments (y-axis). (B) Phenotypic variance of ~1000 segregants (y-axis) within each environment (x-axis). Green colour refers to environments with low phenotypic variance (*Lv*) and pink refers to environments with high phenotypic variance (Hv). The dashed line indicates the median of the distribution. (C) Comparison of phenotypic variance of ~1000 segregants between *Hv* (y-axis) and *Lv* (x-axis) environments. A low regression coefficient indicates poor correlation between the two.

Apart from the genotype, the environments considered also determine the plasticity of an individual. We have previously shown that while a population shows highly buffered phenotype in one environment, this buffering can be lost in others [24]. Hence, we compared the phenotypic variance of the segregants within each environment (Fig 2B). The variance in the 34 environments did not show either a normal or a bimodal distribution but a highly left skewed distribution with a median of 4.2 (Fig 2B). Hence we categorised the environments that were within the first quartile (0 to 8) in the category *Lv* environments. While the remaining 10 environments showed a large range of variance, splitting them into smaller number of environments could have reduced the statistical significance of the variance and slope phenotypes. Therefore, we categorised these 10 environments as *Hv* environments (Fig 2B). We calculated variance of each segregant in *Lv* and *Hv* environments independently, and found no correlation between the two values (Fig 2C). This indicates that a segregant with highly variable phenotype in *Lv* environments can be either plastic or canalised in the *Hv* environments and vice versa. We also calculated mean of segregants across *Hv* and *Lv* environments, and found it to be poorly correlated (*R^2^* = 0.03, Fig S2A). Furthermore, if genetic regulation between random sets of *Lv* environments was as diverse as that between *Hv* and *Lv* environments, then we should observe poor correlation among *Lv* environments. We sampled two random sets of 10 environments each from the *Lv* category and computed correlation of mean values of segregants. These two sets had non-overlapping environments such that the presence of common environments does not bias the correlation. We observed a significantly high correlation between mean across these two sets (*R^2^* = 0.38, *P* < 0.01, Fig S2B), which indicated similar genetic regulation in *Lv* environments, but differential regulation across the *Hv* and *Lv* environments.

### Different loci are pleiotropic in high and low variance environments

Studies have shown that while most yeast growth QTL tends to be environment specific, some loci have pleiotropic effects. A pleiotropic locus is one that has an effect on the phenotype across multiple environments. In order to determine whether plasticity QTL are the same as or a subset of or entirely different from pleiotropic QTL, we carried out QTL mapping in each environment (see Methods). A complete overlap of the large effect QTL and a high overlap of small effect QTL was observed between this study and the original study by Bloom et al. [13] (S1 Table) reconfirming our mapping results. We first compared the pleiotropic loci identified in multiple environments. A locus was designated as pleiotropic if it has an effect in 4 or more environments with a LOD peak within 40kb interval in these environments. Multiple QTL were identified to be pleiotropic across the 34 environments (Table 1).

**Table 1:**
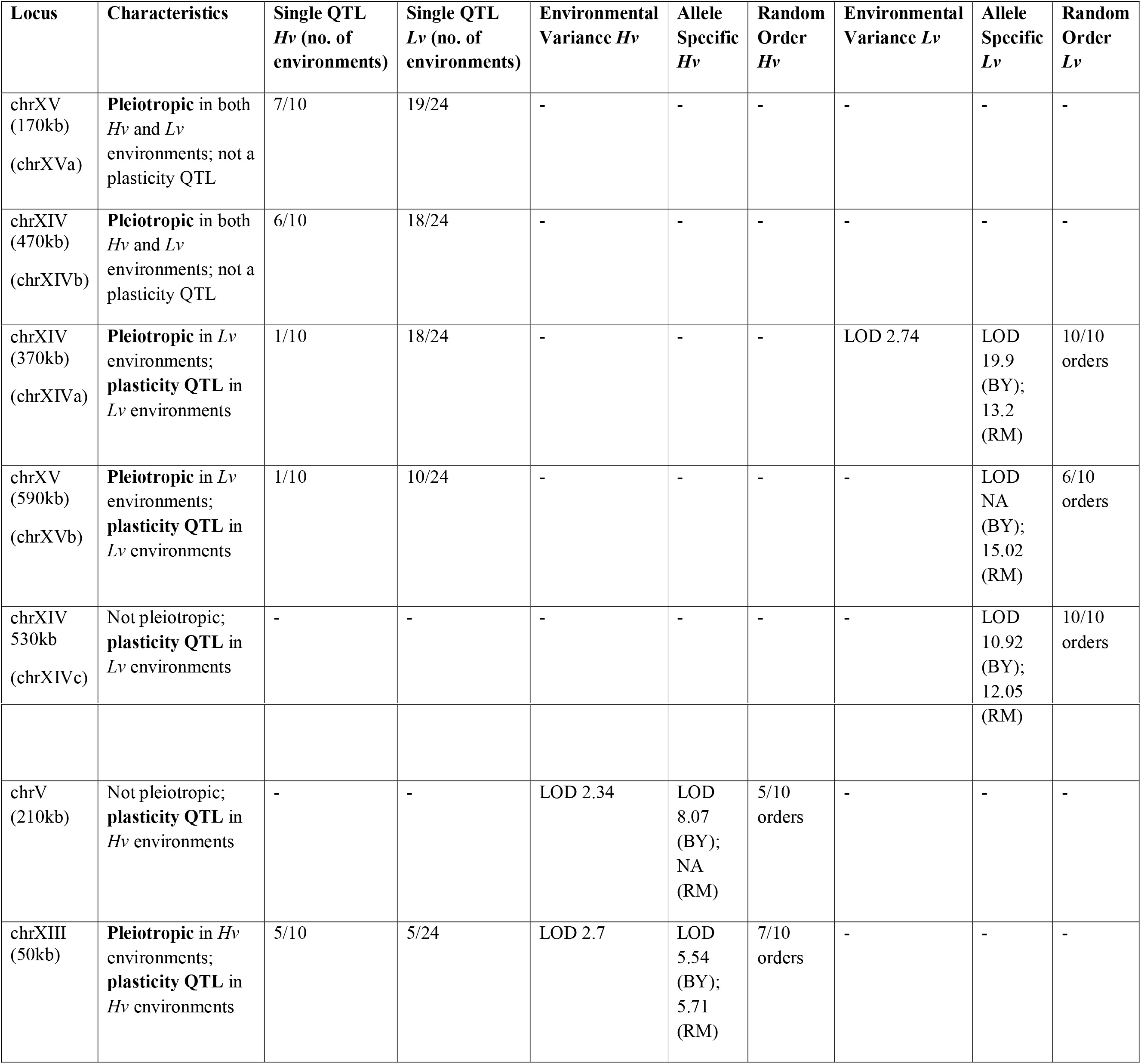
Comparison of QTL and plasticity QTL

We next compared if the pleiotropic loci were different between the *Hv* and *Lv* environments. We found that some pleiotropic loci were common, but others were specific to only *Hv* or *Lv* environments (Fisher’s Exact test *P* < 0.1, Table 1, S1 Table). This shows that there exists a difference in genetic regulation of the phenotype between the *Hv* and *Lv* environments, as predicted by poor correlation of mean across *Hv* and *Lv* environments but strong correlation among *Lv* environments (Fig S2). Previously done fine mapping studies done using the BYxRM segregant populations provide potential candidate genes in many of these loci. Previously, chrXIVb and chrXVa peaks have been identified in multiple environments and fine-mapped to pleiotropic genes like *MKT1* [12] and *IRA2* [12,25] respectively, however in this study neither of these were identified as plasticity QTL in either category of environments. Another pleiotropic QTL, chrXIII locus has been previously associated with yeast chronological lifespan and telomere length with gene *BUL2* as causative [26]. Finally, chrV QTL effected colony morphology with *GPA2* as causal gene [27]. While chrXIVa QTL has not been fine-mapped to any gene, various peaks identified in single QTL and plasticity QTL mapping (see below) indicated that causal gene could be *KRE33*, a protein required for biogenesis of small ribosomal subunit with its human homolog implicated in several types of cancer and premature ageing [28].

### Identifying plasticity QTL using environmental variance

In order to identify plasticity QTL, the first step is to determine a parameter that captures plasticity of segregants. We used modifications of two commonly used parameters: variance and reaction norm or slope [17,29]. Commonly applied data normalisation across environments enhances the power of comparing effect of loci across two environments and helps identifying GEI. However, it also makes the allelic effects symmetric thereby making both alleles equally plastic which results in an inability to distinguish between plastic and canalised alleles (Fig 1A). Therefore, since the aim of this paper was to identify plasticity QTL and not GEI, we normalised the phenotype within an environment but not across environments. While this reduced the power of identifying QTL, the ability to identify plasticity QTL was preserved. Whether one does across-environment normalisation or not, this has no bearing on the QTL identified within an environment [16].

Environmental variance (*Var_E_*) refers to the variance of the phenotype of a segregant across multiple environments. As discussed above, high variance would indicate that the segregant has diverse or plastic phenotype across environments and low variance would suggest that the segregant shows similar phenotype, or canalised behaviour, across environments. Since the scale of variance was different for *Hv* and *Lv* environments (Fig 2B), *Var_E_* was calculated for each segregant independently for each class of environments. As a result, we got two phenotypes for each segregant: *Var_E_* in *Hv* and *Var_E_* in *Lv* environments. We observed no correlation between average phenotype and segregant *Var_E_*, indicating that the two properties were not significantly related (Pearson correlation *P* > 0.1). We then performed QTL mapping for these two phenotypes. While the overall LOD scores identified were lower than conventional single environment QTL mapping, the peaks were significant (Fig 3A, 3D, S2 Table, permutation *P* < 0.01). Two peaks were identified in *Hv* (Fig 3B, 3C) and one in *Lv* environments (Fig 3E) with a LOD score > 2.0 (*P* < 0.01). The highest peak in *Lv* environments, chrXIVa locus was pleiotropic and was unique to this class of environments (Table 1). One peak in *Hv* environments were pleiotropic (chrXIII locus) or whereas the other was not (chrV locus). Interestingly, for both the peaks in *Hv* environments, on chrV, chrXIII, the RM allele had higher environmental variance than BY allele; whereas for the one peak in chrXIVa locus in *Lv* environments, the BY allele showed higher environmental variance (Fig 3, S2 Table). Surprisingly in single QTL mapping, BY allele of chrXIVa, which is the more plastic allele, had lower mean than the RM allele in almost all cases.

**Fig 3:**
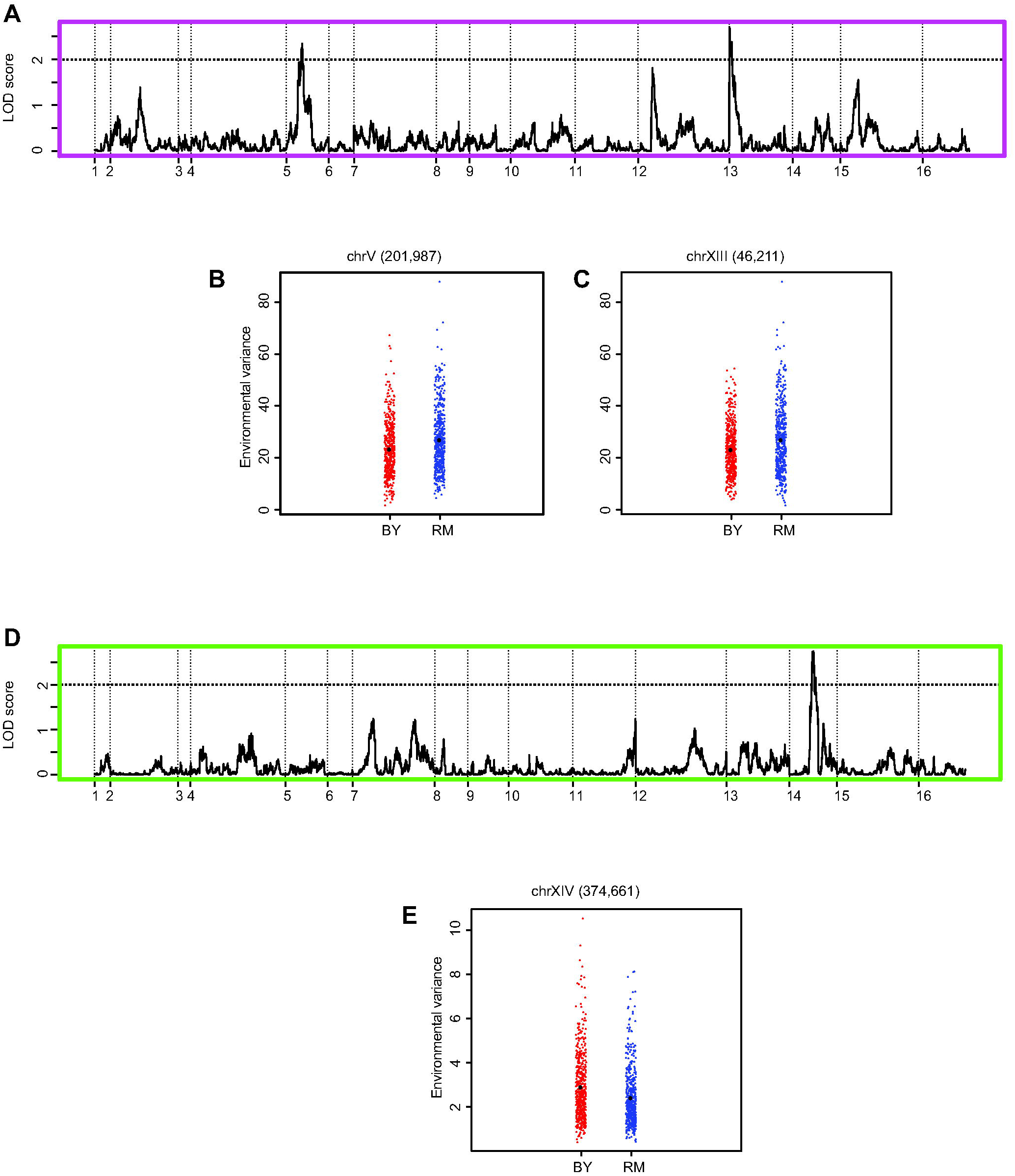
QTL mapping of environmental variance in *Hv* and *Lv* environments. (A) LOD score distribution plot of environmental variance across *Hv* environments. The dashed line represent the LOD cut off of 2.0, permutation *P* < 0.01 (B) Dot plot of marker at chrV (201,987). (C) Dot plot of marker at chrXIII (46,211). (D) LOD score distribution plot of environmental variance across *Lv* environments. The dashed line represent the LOD cut off of 2.0, permutation *P* < 0.01 (E) Dot plot of marker at chrXIV (374,661). Red and blue colours denote BY and RM alleles respectively.

While the environments with variance greater than 8 were categorised as *Hv* environments, as the Fig 2B shows, the highest variable environments show large variance values and can possibly themselves be split further into two subgroups. Therefore, we split 7 *Hv* environments (variance greater than 20) into two subgroups - *Hv_subgroup1* and *Hv_subgroup2* (S1 Table). *Var_E_* was calculated for each segregant independently for each subgroups and QTL mapping was performed as previously discussed (Fig S3, S2 Table). While some loci vary between different subgroups, the large effect chrXIII locus, which was both pleiotropic and plastic in all *Hv* environments, was also identified in both the subgroups (Fig S3, S2 Table) supporting to the original categorisation of *Hv* and *Lv* environments.

Many loci that were pleiotropic across different environments were not identified as plasticity QTL. A stark example is the chrXIVb locus that has been identified as a pleiotropic locus in many environments but had no effect on phenotypic plasticity (Table 1).

### Identifying plasticity QTL using sum of slopes

While *Var_E_* provides an unbiased measure of phenotypic plasticity, it is not sensitive to relatively small changes in the phenotype (Fig 1D). As a result, most GEI studies calculate reaction norms or slopes to identify small effect but significant changes in the phenotype across environments. Usually GEI analysis is performed for a pair of environments [16,21]. As shown by these studies, the steeper the slope of the reaction norm, the more plastic is the genotype. While sensitive, this method can be used only for 4-5 environments or continuums of environments. Large number of heterogeneous environments results in multiple pairwise comparisons that are difficult to both compute and compare. We overcame this shortcoming by computing a novel parameter called sum of slopes (∑*Slope*, see Methods, Fig 1). Briefly, we arrange the environments in different orders and calculate slopes between consecutive environments. The sum of absolute values of these slopes, so that slopes in opposite direction do not cancel each other, gives the value of the parameter. Higher the sum of slope value, more plastic is the individual. Unlike *Var_E_*, sum of slopes will depend upon the order of the environments considered (Fig 1C, 1D). We asked the following questions: how much overlap will be observed in the plasticity QTL mapped using these two different parameters? Will identification of plasticity QTL using sum of slopes depend on the order of the environments?

As done for *Var_E_*, we calculated sum of slopes for each segregant separately for the *Hv* and *Lv* environments. For each category, we used two different strategies to compute the order of the environments. First strategy was to generate random orders, where using permutations, we computed 10 random orders of the environments and then calculated sum of slopes for each segregant for an order and used this as a phenotype for mapping. As a result, we obtained plasticity QTL for each order of the environments, for both *Hv* and *Lv* environments separately (S3 Table, permutation *P* < 0.01). Second strategy was to generate allele specific environmental orders, which takes into consideration that different alleles might have evolved as a result of different selection pressures and hence show canalisation across different orders of environments. While 10 combinations is a substantial number, it may not be exhaustive enough to identify canalisation orders for all alleles. Therefore, we ordered the environments for each allele of each marker independently. For both *Hv* and *Lv* environments independently, for each locus, the environments were ordered to have the least possible sum of slopes for one allele. This order was then used to calculate sum of slopes for all segregants and the values were used for plasticity QTL mapping. The same was done for the other allele separately. Therefore, the total number of environmental orders tested was equal to the product of number of markers, two categories of environment and two alleles. Thus, the QTL were mapped for a canalised mean of each allele for each locus, in both categories of environments (Fig 4, S4 Table, permutation *P* < 0.01).

**Fig 4:**
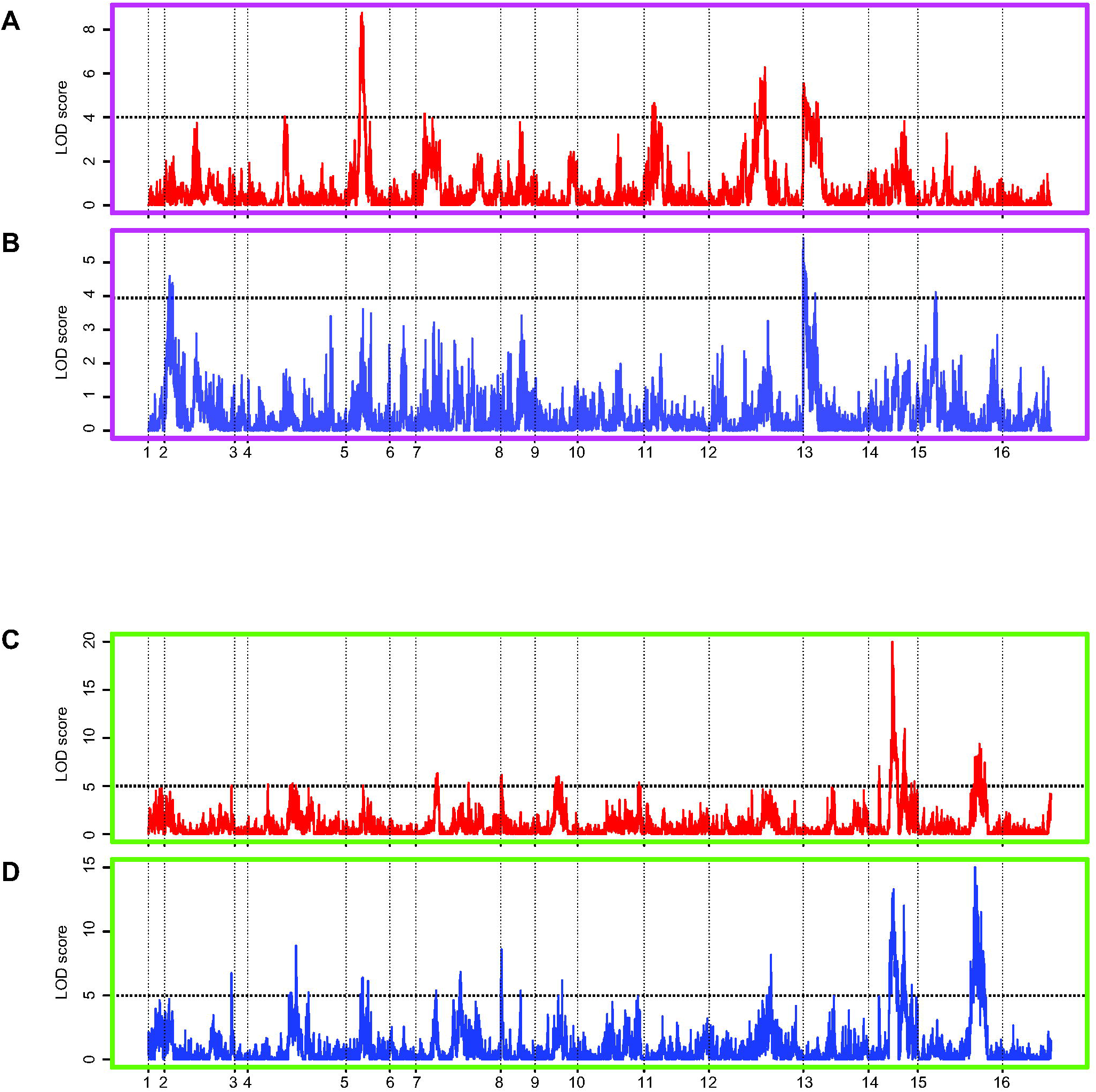
QTL mapping of reaction norms in *Hv* and *Lv* environments using allele specific orders. (A) and (B) show LOD score distribution plots of reaction norms using allele specific order across *Hv* environments. The dashed line represent the LOD cut off of 4.0 in A and B respectively, permutation *P* < 0.01. (C) and (D) show LOD score distribution plots of reaction norms using allele specific order across *Lv* environments. The dashed line represent the LOD cut off of 5.0 in C and D respectively, permutation *P* < 0.01. Red and blue plots indicated QTL mapping performed by canalising BY and RM alleles, respectively.

Higher LOD scores and larger number of plasticity QTL were identified for sum of slopes than that were identified for environmental variance (Table 1, S2, S3 Table). For random order analyses, the plasticity QTL identified depended on the order of the environments. We compiled the results to identify peaks that were identified in most environmental orders. Certain plasticity QTL were identified in more than half of 10 random environmental orders, i.e. they were independent of the environmental order (Table 1). While 4 peaks were identified in majority of the environmental orders consisting of *Lv* environments, only a single peak was consistently identified in *Hv* environments (Table 1, S3 Table). These loci included the ones identified using *Var_E_*, as well as unique to sum of slopes (Table 1). The loci were identified with higher LOD scores in the *Lv* than the *Hv* environments (S3 Table).

As noted in random order analyses, higher LOD scores and more peaks were identified using sum of slopes than *Var_E_* (Fig 4). Distinct sets of peaks were identified in *Hv* and *Lv* environments using allele specific environmental orders (S4 Table). Additionally, like the plasticity QTL identified depended on the random order, the identification of the plasticity QTL using allele specific order depended on the allele whose mean was canalised (Fig 4, S4 Table). However, we also identified plasticity QTL that were independent of the allele whose mean effect was canalised, i.e. they were identified independent of whether the RM or BY allele was canalised. These overlapped with the plasticity QTL that were identified in most random orders of environments (Table 1).

We compared plasticity QTL identified using three strategies: *Var_E_*, sum of slopes with random orders and sum of slopes with allele specific orders (Table 1). As proposed in Fig 1, both *Var_E_* and sum of slopes are capable of identifying differences in plasticity to different extents and measuring both of them is required to identify the genetic regulators of phenotypic plasticity. While several QTL were specific to the parameter or environmental order used, two loci chrXIII in *Hv* and chrXIVa in *Lv* environments were identified in all three methods (Table 1). Identification of these plasticity QTL through independent strategies emphasises their definite ability to regulate phenotypic plasticity.

Comparison of sum of slopes revealed that, as expected, the value of this parameter was less for *Lv* than for *Hv* environments. However, canalisation of mean of the allele, i.e. the lowest sum of slopes of mean, as done for allele specific order, did not necessarily result in reduced sum of slopes of the segregants carrying the allele (S4 Table). For plasticity QTL that were identified independent of the allele, the same allele had higher sum of slopes of segregants independent of the allele whose mean was canalised (S3, S4 Table). This explains why some plasticity QTL were identified irrespective of the environmental order. Furthermore, this shows that canalisation of the population mean does not always reflect canalisation of the individuals within the population (Fig 5A, 5B). An allele can have a canalised mean but differential plasticity of individuals. This was observed for both *Hv* and *Lv* environments (Fig 5A, 5B). Furthermore, our results show that while environmental order can uncover the difference in plasticity between two alleles, a canalised allele will always be canalised independent of the environmental order (S4 Table).

**Fig 5:**
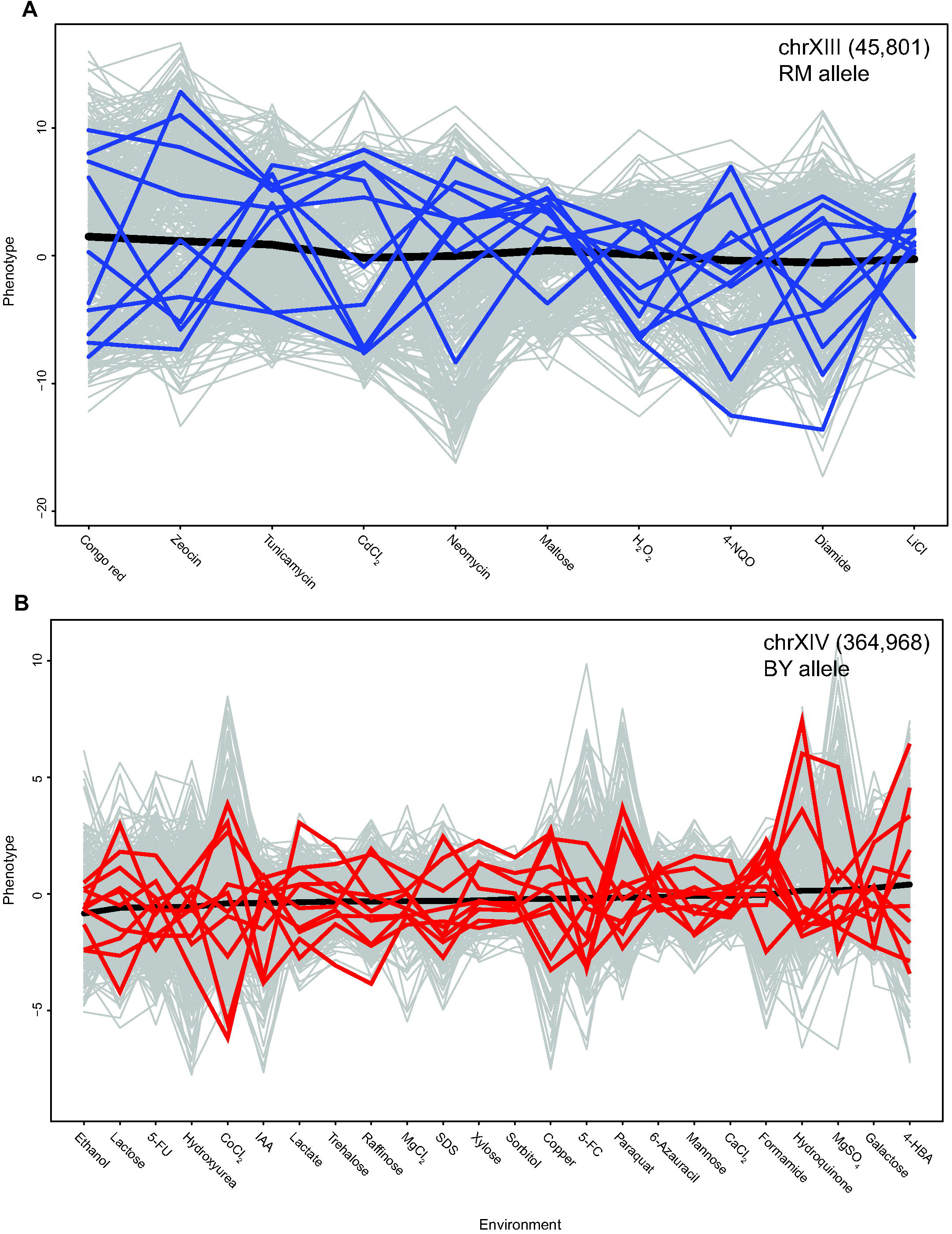
Phenotypic plasticity observed within canalised mean effects. Reaction norms of segregants carrying RM allele of marker chrXIII (45,801) in *Hv* environments (A), and BY allele of marker chrXIV (364,968) in *Lv* environments (B). In both the plots, the environments are arranged such that the mean phenotype, denoted by the black line, has the least possible value of sum of slopes. Reaction norms for 10 random segregants have been highlighted as blue, RM, and red, BY in the two plots and reaction norms of other segregants are represented in grey lines.

High variance of sum of slopes within an allele would indicate diversity of phenotypic plasticity. While there was no association between mean and variance of segregant values across environments, we found that there was a positive association between the mean and variance of sum of slopes between various alleles in both *Hv* and *Lv* environments indicating that the allele with higher sum of slopes also showed more diversity (Fig 6). Therefore, the segregants carrying the more plastic allele did not show same pattern of phenotypic plasticity but demonstrated a diversity of patterns, potentially to facilitate adaptation to diverse environments. Hence, our results show that the more plastic allele also results in revelation of hidden reaction norms.

**Fig 6:**
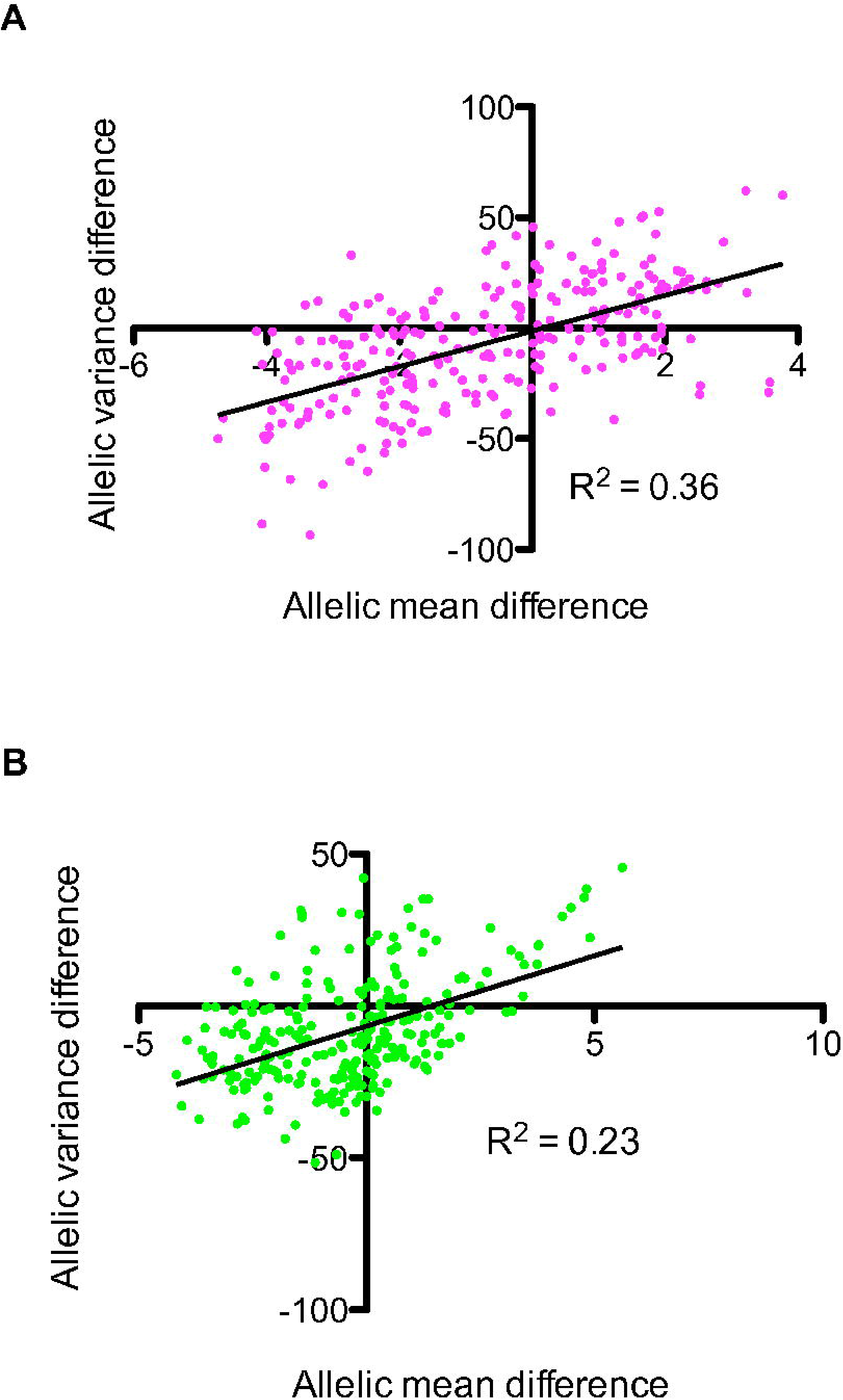
Comparison of mean and variance of allelic reaction norms. Comparison of difference in mean and variance of the alleles of peaks identified in 10 different random orders in *Hv* (A) and *Lv* (B) environments. x-axis shows the difference between mean value of sum of slopes of alleles for different peaks, BY-RM, and y-axis refers to difference between variance of sum of slopes of alleles, BY-RM. See S3 Table for more details.

## DISCUSSION

Our study identifies loci with differential effects on phenotypic plasticity in heterogeneous environments. We show that regulation of phenotypic plasticity is overlapping but different than the regulation of phenotypic variation in each environment. This has implications not only on adaptation and evolution, but also on understanding the genetic architecture of genotype-phenotype map. While different plasticity QTL were identified using different parameters of plasticity and in different environmental orders, some of these plasticity QTL were identified in all mapping methods indicating their robust role in regulating phenotypic plasticity.

Phenotypic plasticity is a property of the genotype, unveiled by the environments. We show that environments can be divided into two categories based on phenotypic variance of the population and *Hv* and *Lv* environments (Fig 2). Such a distinction has been hypothesised by previous studies [4], which propose that when a population is adapted to a particular environment, then stabilising selection acts on the population, such that most individuals of the population show similar phenotype which is close to the fitness optimum (low variance). When the population encounters a novel or rare environment, this buffering is perturbed releasing high diversity of individual phenotypes (high variance), which can facilitate adaptation. In the light of this current evolutionary understanding of plasticity and canalisation, we infer our results from a biparental population as follows: the *Lv* environments are the ones in which either one or both strains have adapted to in the course of their evolutionary history whereas the *Hv* environments are potentially novel environments [30]. This conclusion is further facilitated by identification of different QTL as well plasticity QTL in both these categories of environments (Table 1). Differential enrichment of pleiotropic QTL indicates a common regulation of the phenotype in the canalised or *Lv* environments. Additionally, disruption of canalisation in the recombinant population may explain why the large effect and consistent plasticity QTL were identified in *Lv* than the *Hv* environments. Genetic recombination disrupts the evolved canalisation mechanisms therefore resulting in identification of plasticity QTL in *Lv* environments, whereas poor or no canalisation mechanisms exist for *Hv* environments, which results in high plasticity of all alleles. This results in reducing the LOD score of plasticity QTL identified.

As proposed in Fig 1, our results show that plasticity QTL are not same as pleiotropic QTL. Almost all loci show GEI and large effect pleiotropic loci show large effect GEI [16]. However, we observed only a partial overlap between pleiotropic QTL and plasticity QTL. While some large effect QTL (like chrXIII and chrXIVa) also had pleiotropic effects, others like chrV and chrXIVc did not show pleiotropy but were equally significant plasticity QTL. In fact, while chrXIVa and chrXIVc were in a relative close physical distance, within 160kb (Table 1), they had opposite effects on plasticity of the alleles: BY allele of chrXIVa showed high plasticity and RM allele of chrXIVc showed high plasticity (S3, S4 Table). This indicates that genetic regulation of phenotypic plasticity is overlapping, but different than genetic regulation within each environment. This further emphasises that in order to understand the genotype-phenotype map and the function of identified molecular regulatory hubs, it is important to not only understand their effects in one environment or phenotypes but across different environments.

In a previous study, we showed the biological implication of mean and variance of a population [24]. We showed that higher variance was associated with phenotypic manifestation of cryptic or hidden variants. Additionally, a high phenotypic variance could either be associated with a higher or a lower phenotypic mean depending on the environment. Here we show a strong correlation between mean and variance of phenotypic plasticity (Fig 6). Interestingly, in both *Hv* and *Lv* environments, the allele with a higher mean of plasticity also had a higher variance (Fig 6). This indicates that segregants containing the more plastic alleles exhibit a diverse range of phenotypic plasticity, potentially to facilitate adaptation in diverse environmental conditions. The high variance of plasticity values (both *Var_E_* and sum of slopes) suggests epistasis resulting in revelation of hidden reaction norms [4] or cryptic genetic variants with diverse effects across environments. Along with shedding light on mechanisms of regulation of phenotypic plasticity, this suggests an association between genetic regulation of cryptic genetic variation and phenotypic plasticity [31].

In conclusion, by identifying genetic regulators of phenotypic plasticity and canalisation, our results highlight that genetic regulation of a phenotype in an environment may depend not only upon mechanisms directly evolved in that environment but maybe a result of evolution in a diverse range of environments [15,32]. While commenting on the evolutionary nature of the identified plasticity QTL is beyond the scope of our results, our study opens new avenues of exploring population genetic data and understanding the underlying basis of the genetic architecture. Differential regulation of phenotypic plasticity provides a potential reason underlying the high interconnectivity observed in the genotype-phenotype map. This interconnectivity could be an outcome of cross talk between different genetic modules that either maintain canalisation or induce plasticity across different environments and phenotypes. This has profound implications, especially on understanding adaptation mechanisms in naturally occurring plant and animal populations, development [33] as well as understanding the molecular basis of regulation of complex human diseases highly susceptible to environmental conditions [34] such as metabolic and psychological disorders.

## SUPPLEMENTARY TABLE LEGENDS

**S1 Table: Comparison of QTL identified in each environment independently**

**S2 Table: Plasticity QTL identified using environmental variance (*Var_E_*) in *Hv* and *Lv* environments**

**S3 Table: Plasticity QTL identified using sum of slopes in 10 randomly generated orders of environment in *Hv* and *Lv* environments**

**S4 Table: Plasticity QTL identified using sum of slopes in allele specific orders of environment in *Hv* and *Lv* environments**

## SUPPLEMENTARY FIGURE LEGENDS

**Figure S1: Normal distribution of environmental variance (*Var_E_*) phenotype**. (A) Histogram showing the normal distribution of environmental variance across all environments. x-axis shows classes of variance with an interval size of *Var_E_* = 1.0 and y-axis shows the number of segregants showing a particular variance value. (B) QQ plot comparing the observed variance of segregants with the expected variance, given the distribution in normal. x-axis shows the expected value of a distribution of 1007 individuals with a mean of 9.48 and standard deviation of 3.46 (as observed in the current distribution) and y-axis shows the observed values of the segregants. (C) Histogram showing the normal distribution of environmental variance across *Lv* environments. x-axis shows classes of variance with an interval size of *Var_E_* ranging from 0.25 to 0.5, and y-axis shows the number of segregants showing a particular variance value. (D) Histogram showing the normal distribution of environmental variance across *Hv* environments. x-axis shows classes of variance with an interval size of *Var_E_* = 2.0 and y-axis shows the number of segregants showing a particular variance value.

**Figure S2: Comparison of mean of segregants across different groups of environments**. (A) Comparison of mean values of all segregants across 24 *Hv* environments (x-axis) with that across *Lv* environments (y-axis). (B) Comparison of mean values of all segregants across two mutually exclusive sets of 10 environments each, chosen from the 24 *Lv* environments, set 1 (x-axis) and set 2 (y-axis).

**Figure S3: LOD score distribution plot of environmental variance in Hv_subgroup1 and Hv_subgroup2**. The dashed line represent the LOD cut off of 1.0, permutation *P* < 0.05.

